# Hidden diversity and macroevolutionary mode of *Caulimoviridae* uncovered by euphyllophyte paleoviruses

**DOI:** 10.1101/170415

**Authors:** Zhen Gong, Guan-Zhu Han

## Abstract

Few viruses have been documented in plants outside angiosperms. Endogenous viral elements (paleoviruses) provide ‘molecular fossils’ for studying the deep history and macroevolution of viruses. Endogenous plant pararetroviruses (EPRVs) are widespread across angiosperms, but little is known about EPRVs in earlier branching plants. Here we use a large-scale phylogenomic approach to investigate the diversity and macroevolution of plant pararetroviruses (formally known as *Caulimoviridae*). We uncover an unprecedented and unappreciated diversity of EPRVs in the genomes of gymnosperms and ferns. The known angiosperm viruses only constitute a minor part of the *Caulimoviridae* diversity. By characterizing the distribution of EPRVs, we show that no major euphyllophyte lineages escape the activity of *Caulimoviridae*, raising the possibility that many exogenous *Caulimoviridae* remain to be discovered in euphyllophytes. We find that the copy numbers of EPRVs are generally high, suggesting that EPRVs define a unique group of repetitive elements and represent major components of euphyllophyte genomes. Phylogenetic analyses reveal an ancient monilophyte origin of *Caulimoviridae* and at least three independent origins of *Caulimoviridae* in angiosperms by cross-division transmissions. Our findings uncover the remarkable diversity of *Caulimoviridae* and have important implications in understanding the origin and macroevolution of plant pararetroviruses.

---

Endogenous viral elements (EVEs), the viral sequences integrated into their hosts’ genomes, document past virus (paleovirus) infections and provide ‘molecular fossils’ for studying the deep history of viruses^1^. EVEs lay the foundation of an emerging filed, Paleovirology^1,2^. The best characterized EVEs are endogenous retroviruses (ERVs)^3^. The replication of retroviruses requires integration into their hosts’ genomes. On occasion, retroviruses infect germ lines of their hosts, and the integrated retroviruses, namely ERVs, become vertically inherited. ERVs are widespread and highly abundant in the genomes of vertebrates^3^; for example, ERVs make up 5%-8% of the human genome^4^. Recently, endogenous non-retroviral elements have been increasingly identified by comparative genomic analyses, which reveal the remarkable diversity, deep history, and macroevolution of related viruses^5,6^. EVEs (especially ERVs) were pervasively co-opted for the hosts’ biology, ranging from placentation, to inhibition of exogenous viral infection, to regulation of innate immunity^7-9^.

Like retroviruses, two families of viruses with double-stranded DNA genomes replicate through RNA intermediates, known as pararetroviruses or DNA reverse-transcribing viruses^10^. Unlike retroviruses, these pararetroviruses lack integrase, and thus integration into host genomes is not essential for their replication. Pararetroviruses infect vertebrates (*Hepadnaviridae*) and plants (*Caulimoviridae*). Evolutionary analyses suggest that *Hepadnaviridae* and *Caulimovirida*e originated independently from retrotransposons with long terminal repeats (LTRs)^11^. Endogenous hepadnaviruses have been increasingly identified in the genomes of many species of Aves and reptiles^12-14^. The copy number of endogenous hepadnaviruses in the host genomes is very low (usually around ten copies)^12-14^. The identification of endogenous hepadnaviruses reveals the prevalent nature and deep history (more than 207 million years) of *Hepadnaviridae* in vertebrates^12-15^.

The *Caulimoviridae* family, pararetroviruses infecting plants, is classified into eight genera, namely *Caulimovirus*, *Soymovirus*, *Tungrovirus*, *Badnavirus*, *Solendovirus*, *Cavemovirus*, *Rosadnavirus*, and *Petuvirus*^16^. The genome size of *Caulimoviridae* is usually between 6,000 and 8,000 base pairs (bps), encoding one to eight open reading frames (ORFs). The proteins (or domains) common to *Caulimoviridae* include movement protein (MP), coat protein (CP), aspartic protease (AP or PR), reverse transcriptase (RT), and RNase H1 (RH)^16^. While the replication of *Caulimoviridae* does not require integration into host genomes, endogenous plant pararetroviruses (EPRVs) were identified in many angiosperms at pre-genomic era^17^, for example banana^18^ and tobacco^19^. Genome-scale data provide important resource to explore the distribution and diversity of EPRVs within plant genomes, which would improve our understanding of the macroevolution of *Caulimoviridae* and the relationship between viruses and their hosts. By mining a variety of plant genomes, Geering et al.^20^ identified a novel lineage of EPRVs in flowering plants (angiosperms), which was designated ‘Florendovirus’ and was thought to constitute a new genus within *Caulimoviridae*. However, EPRVs have not been detected in the genomes of plants outside angiosperms^20^.

In this study, we use a large-scale phylogenomic approach to investigate whether EPRVs are present in the genomes of plants outside angiosperms. By mining ten gymnosperm and six fern genomes, we identified EPRVs in the genomes of nearly all these gymnosperms and ferns. Phylogenetic analyses using the newly identified EPRVs together with other angiosperm viruses reveal an unappreciated diversity of *Caulimoviridae* and show that the known angiosperm viruses only constitute a minor part of the *Caulimoviridae* diversity. The newly identified EPRVs in gymnosperms and ferns provide important and novel insights into the diversity, distribution, and macroevolution of *Caulimoviridae*.

## Results

### Identification of EPRVs in gymnosperms and ferns

We used a combined similarity search and phylogenetic analysis approach to screen the genomes of ten gymnosperms, six ferns, and four other earlier branching plant species (*Selaginella moellendorffii*, *Physcomitrella patens, Marchantia polymorpha*, and *Klebsormidium flaccidum*) for the presence of EPRVs (Fig. 1 and Supplemental Table 1). Briefly, similarity search with the protein sequences of representative *Caulimoviridae* was performed against these plant genomes (Fig. 1 and Supplemental Table 1). Given RT and RH of *Caulimoviridae* share significant similarity with retrotransposons and other reverse-transcribing viruses, EPRVs were further identified and confirmed by phylogenetic analyses (see Methods). We found that EPRVs are present in the genomes of nearly all the gymnosperms and ferns investigated in this study (Fig. 1), suggesting that EPRVs are prevalent and widespread in gymnosperms and ferns. EPRVs were not identified in the genome of the fern *Ceratopteris richardii*, which does not necessarily indicate the absence of EPRVs but is more likely due to the low-density coverage (1.082×) of its genome sequencing (only 1% of its genome is covered by the genome assembly)^21^. No EPRV was detected in the genomes of the lycophyte *S. moellendorffii*, the moss *P. patens*, the liverwort *M. polymorpha*, and the charophyte *K. flaccidum* (Fig. 1). Together with previous reports of EPRVs in angiosperms^17-20^, we conclude that EPRVs are widespread in the genomes of euphyllophytes (ferns and seed plants).

**Figure 1.**
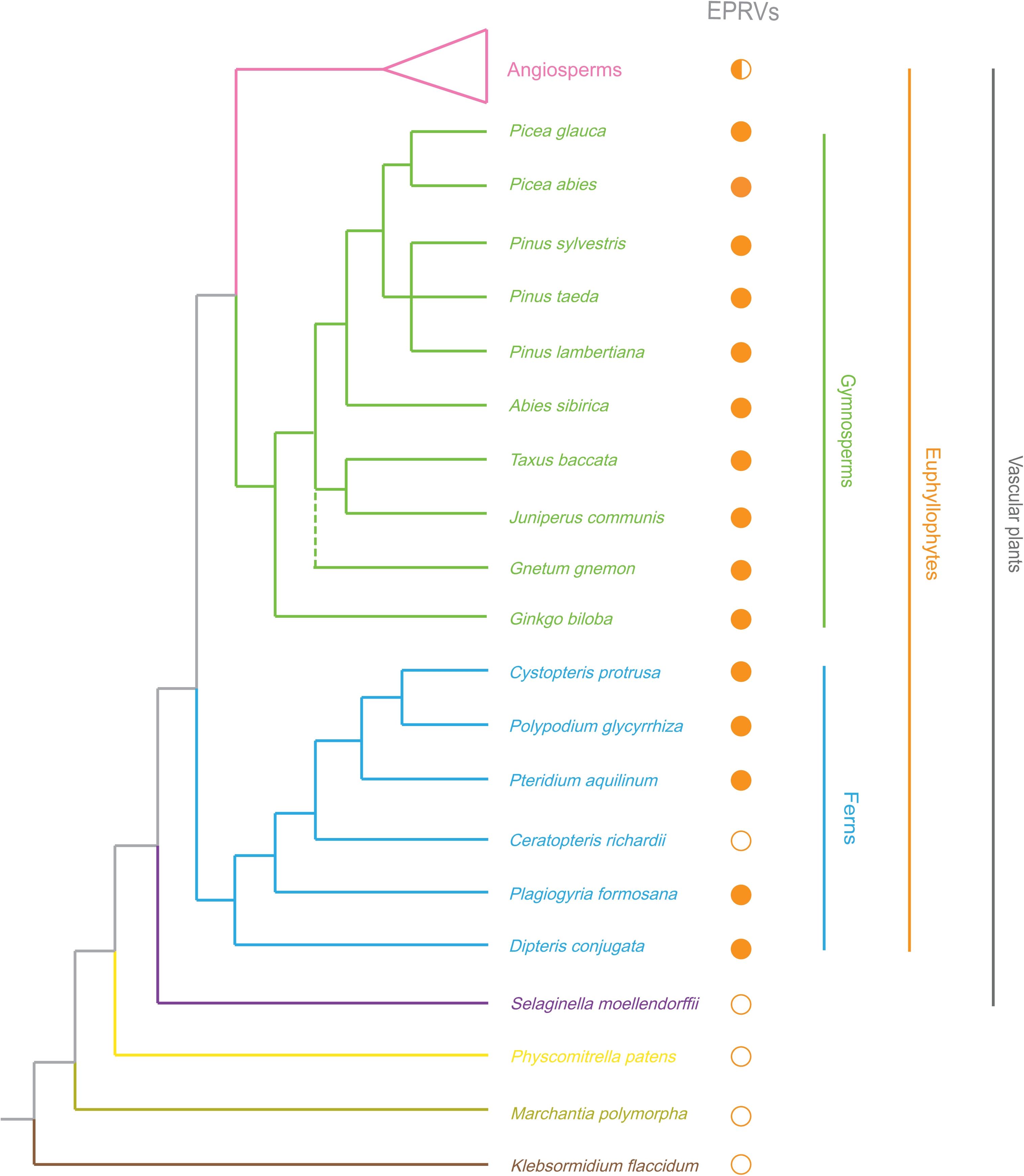
Distribution of EPRVs within plant genomes. The phylogenetic relationship of plant species is based on Refs. 21, 31, 32, 44-46. Different plant divisions were labelled in different colors, and angiosperms, gymnosperms, and ferns were labelled in pink, green, and blue, respectively. The presence and absence of EPRVs were marked with solid and open circles around the related species. The half-filled circle indicates that EPRVs have been identified in some but not all the angiosperms.

The copy numbers of EPRVs within the genomes of gymnosperms and ferns appear to vary widely across plant species. We estimated that the genomes of the gymnosperms and ferns contain 112 – 20,579 copies of EPRVs (Table 1). However, the genomes of some gymnosperms and ferns are of low coverage, these estimated numbers should be taken with cautions. On the other hand, for the three genomes that >90% are covered by genome assembly (*Pinus taeda, Picea glauca*, and *Ginkgo biloba*), the EPRV copy numbers were estimated to be 6,733, 2,520, and 481, respectively. Our results suggest that EPRVs might represent major components of plant genomes.

**Table 1.**
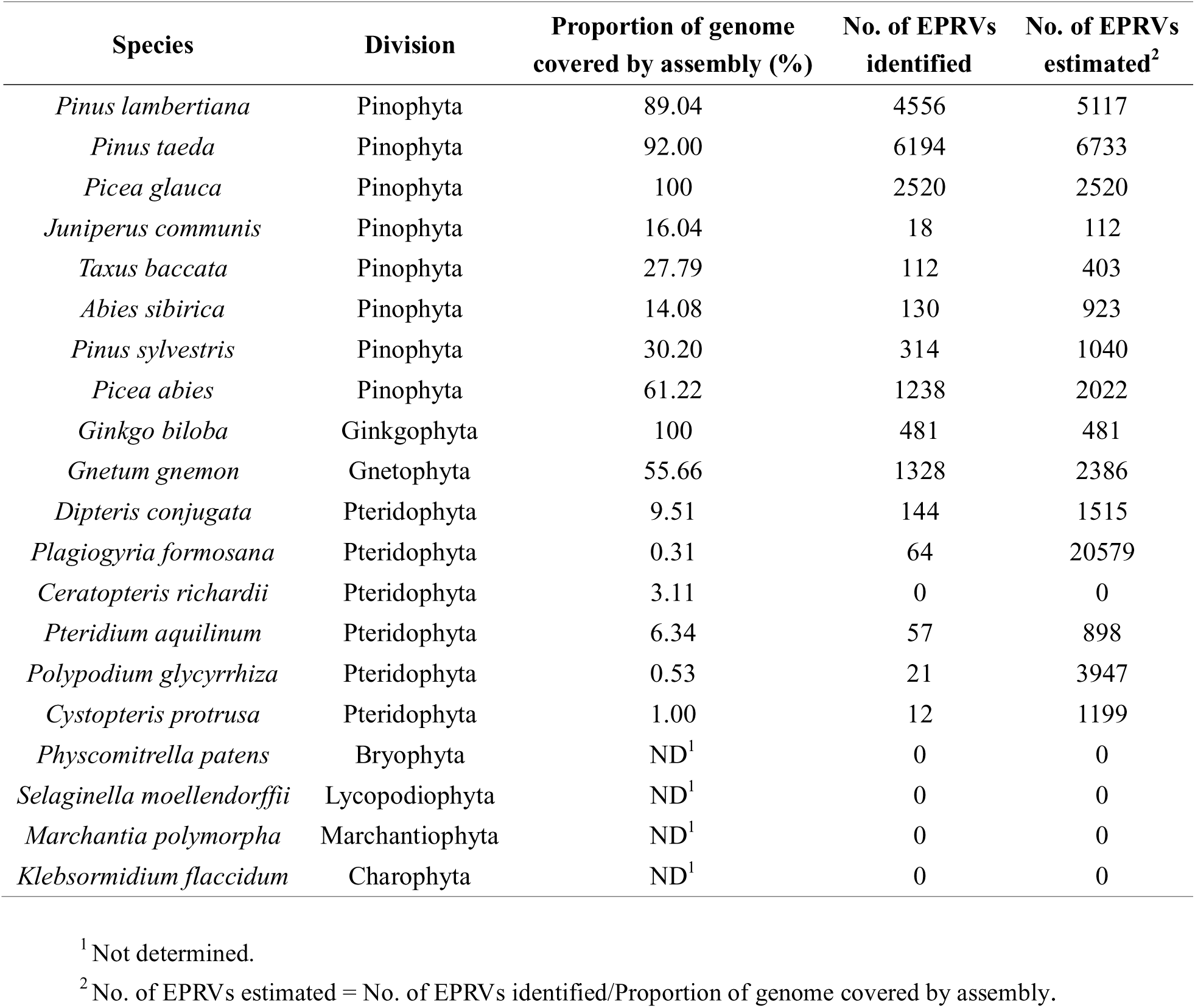
Copy number of EPRVs identified and estimated in plant genomes.

| Species | Division | Proportion of genome covered by assembly (%) | No. of EPRVs identified | No. of EPRVs estimated<sup>2</sup> |
| --- | --- | --- | --- | --- |
| *Pinus lambertiana* | Pinophyta | 89.04 | 4556 | 5117 |
| *Pinus taeda* | Pinophyta | 92.00 | 6194 | 6733 |
| *Picea glauca* | Pinophyta | 100 | 2520 | 2520 |
| *Juniperus communis* | Pinophyta | 16.04 | 18 | 112 |
| *Taxus baccata* | Pinophyta | 27.79 | 112 | 403 |
| *Abies sibirica* | Pinophyta | 14.08 | 130 | 923 |
| *Pinus sylvestris* | Pinophyta | 30.20 | 314 | 1040 |
| *Picea abies* | Pinophyta | 61.22 | 1238 | 2022 |
| *Ginkgo biloba* | Ginkgophyta | 100 | 481 | 481 |
| *Gnetum gnemon* | Gnetophyta | 55.66 | 1328 | 2386 |
| *Dipteris conjugata* | Pteridophyta | 9.51 | 144 | 1515 |
| *Plagiogyria formosana* | Pteridophyta | 0.31 | 64 | 20579 |
| *Ceratopteris richardii* | Pteridophyta | 3.11 | 0 | 0 |
| *Pteridium aquilinum* | Pteridophyta | 6.34 | 57 | 898 |
| *Polypodium glycyrrhiza* | Pteridophyta | 0.53 | 21 | 3947 |
| *Cystopteris protrusa* | Pteridophyta | 1.00 | 12 | 1199 |
| *Physcomitrella patens* | Bryophyta | ND<sup>1</sup> | 0 | 0 |
| *Selaginella moellendorffii* | Lycopodiophyta | ND<sup>1</sup> | 0 | 0 |
| *Marchantia polymorpha* | Marchantiophyta | ND<sup>1</sup> | 0 | 0 |
| *Klebsormidium flaccidum* | Charophyta | ND<sup>1</sup> | 0 | 0 |

<sup>1</sup> Not determined.
<sup>2</sup> No. of EPRVs estimated = No. of EPRVs identified/Proportion of genome covered by assembly.

### Diversity and classification of *Caulimoviridae*

To explore the relationship between the newly identified gymnosperm and fern EPRVs and the known angiosperm *Caulimoviridae*, we inferred a phylogeny of representative exogenous and endogenous viruses of *Caulimoviridae* using the highly conserved RT-RH proteins with the retrotransposon *Ty3* as the outgroup. Our phylogenetic analysis reveals an extraordinarily large diversity of the *Caulimoviridae* family, which has never been appreciated previously (Fig. 2). The known eight viral genera and florendoviruses fell well within the diversity of EPRVs of gymnosperms and ferns. It follows that the previously known angiosperm *Caulimoviridae* constitute only a minor part of its diversity.

**Figure 2.**
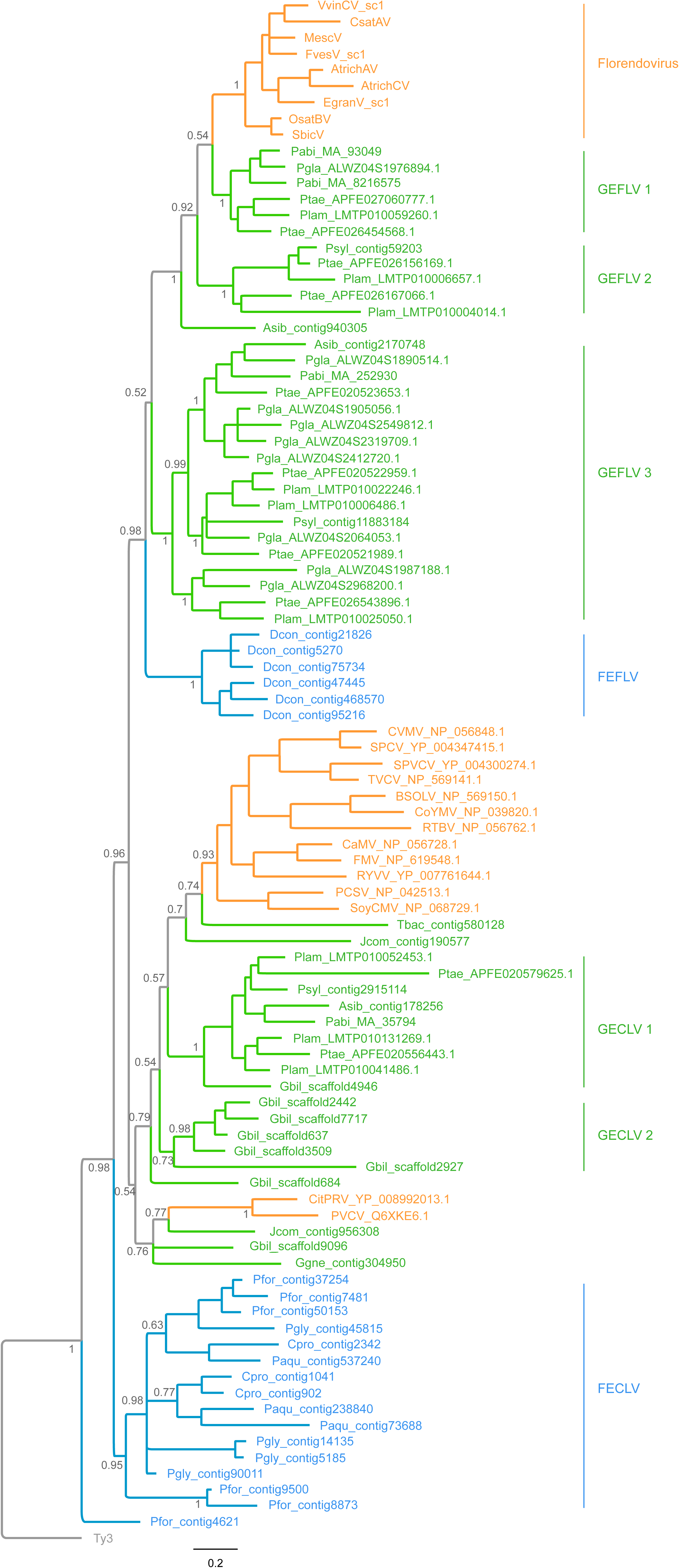
Phylogenetic relationship of representative exogenous and endogenous *Caulimoviridae*. The phylogenetic tree was inferred based on the RT-RH protein sequences using a Bayesian method. The tree was rooted using the *Ty3* LTR retrotransposon as the outgroup. Bayesian posterior probabilities were shown on the selected nodes. *Caulimoviridae* of angiosperms, gymnosperms, and ferns were highlighted in orange, green, and blue, respectively. Virus abbreviations: CVMV, Cassava vein mosaic virus; SPCV, Sweet potato caulimo-like virus; SPVCV, Sweet potato vein clearing virus; TVCV, Tobacco vein clearing virus; BSOLV, Banana streak OL virus; CoYMV, Commelina yellow mottle virus; RTBV, Rice tungro bacilliform virus; CaMV, Cauliflower mosaic virus; FMV, Figwort mosaic virus; RYVV, Rose yellow vein virus; PCSV, Peanut chlorotic streak virus; SoyCMV, Soybean chlorotic mottle virus; CitPRV, Citrus endogenous pararetrovirus; PVCV, Petunia vein clearing virus; VvinCV_sc1, *Vitis vinifera* C virus sequence cluster 1; CsatAV, *Cucumis sativus* A virus; MescV, *Manihot esculenta* virus; FvesV_sc1, *Fragaria vesca* virus sequence cluster 1; AtrichAV, *Amborella trichopoda* A virus; AtrichCV, *Amborella trichopoda* C virus; EgranV_sc1, *Eucalyptus grandis* virus sequence cluster 1; OsatBV, *Oryza sativa* B virus; SbicV, *Sorghum bicolor* virus. Species abbrevations: Pabi, *Picea abies*; Pgla, *Picea glauca*; Ptae, *Pinus taeda*; Plam, *Pinus lambertiana*; Psyl, *Pinus sylvestris*; Asib, *Abies sibirica*; Gbil, *Ginkgo biloba*; Jcom, *Juniperus communis*; Ggne, *Gnetum gnemon*; Tbac, *Taxus baccata*; Pfor, *Plagiogyria formosana*; Pgly, *Polypodium glycyrrhiza*; Cpro, *Cystopteris protrusa*; Paqu, *Pteridium aquilinum*; Pgly, *Polypodium glycyrrhiza*.

Our phylogenetic analysis identified at least seven monophyletic groups of EPRVs with high supports (Bayesian posterior probability > 0.95) in gymnosperms and ferns. These clades were designated gymnosperm endogenous florendovirus-like virus 1-3 (GEFLV 1-3), fern endogenous florendovirus-like virus (FEFLV), gymnosperm endogenous caulimovirus-like virus 1-2 (GECLV 1-2), and fern endogenous caulimovirus-like virus (FECLV), respectively (Fig. 2). The host of each clade is restricted to one plant division (except GECLV 1) (Fig. 2). The divergence within one of these clades is comparable to and even greater than that of one known *Caulimoviridae* genus. Some gymnosperm and fern EPRVs are not readily classified, either because one virus formed one branch or because the branches are not strongly supported.

### Macroevolutionary mode of *Caulimoviridae*

To estimate the relative importance of co-speciation and host switching in the macroevolution of *Caulimoviridae*, we performed a global assessment of the correspondence between *Caulimoviridae* and host phylogenetic trees using the event-based approach. The analyses found no significant signal for co-speciation (*p* values > 0.05; Table 2), suggesting co-speciation might not play a predominant role in the diversification of *Caulimoviridae*.

**Table 2.** Number of events experienced by virus lineages.

| Event costs<sup>1</sup> | Total cost | Cospeciation<sup>2</sup> | Duplication<sup>2</sup> | Duplication & host switching<sup>2</sup> | Loss<sup>2</sup> | Failure to diverge<sup>2</sup> | P-value<sup>3</sup> | P-value<sup>4</sup> |
| --- | --- | --- | --- | --- | --- | --- | --- | --- |
| -1, 0, 0, 0, 0 | -8 | 8-8 | 2-3 | 13-14 | 7-20 | 0-0 | >0.05 | >0.05 |
| 0, 1, 1, 2, 0 | 20 | 4-4 | 3-5 | 15-17 | 0-0 | 0-0 | >0.05 | >0.05 |

<sup>1</sup>Event costs are for cospeciation, duplication, duplication & host switching, loss, and failure to diverge, respectively.
<sup>2</sup>Number of events is expressed as ranges that result in the same cost.
<sup>3</sup>Random tip mapping method with sample size of 500.
<sup>4</sup>Random parasite tree method with sample size of 500.

Our phylogenetic analysis shows that some newly described fern EPRVs occupy important phylogenetic positions -- basal to all the other known *Caulimoviridae*. Ancestral state reconstruction reveals that the *Caulimoviridae* family originated in ferns (Supplemental Fig. 1). Our phylogenetic analysis also shows that the angiosperm viruses form three independent monophyletic groups: two consist of known eight genera of exogenous viruses and one consists of florendoviruses (Fig. 2). The three angiosperm viral groups are only distantly related to each other. The phylogenetic relationship among euphyllophyte viruses indicate that the angiosperm viruses originated multiple times probably through cross-division transmission from gymnosperms.

### Genome structure evolution of *Caulimoviridae*

To explore the genome structure evolution within the *Caulimoviridae* family, we reconstructed the consensus genome sequences of EPRVs (Supplemental Data 1-5). Given the fern genomes are of low coverage, we only reconstructed the genomes of gymnosperm EPRVs, and one representative for each of the five gymnosperm EPRV clades were inferred. These gymnosperm EPRV genomes vary wildly in size (from 6,061 to 8,109 bps) and ORF organization (Fig. 3 and Supplementary Data). Conserved Domain (CD) searches show that protein domains common to all the EPRVs include MP, AP, RT, and RH, suggesting that the gymnosperm EPRVs exhibit a similar protein architecture as the angiosperm *Caulimoviridae* (Fig. 3). No CP homologs were identified in the consensus genome sequences, possibly due to the rapid nature of its evolution. However, we identified the zinc-finger CCHC motif, a hallmark of the CP protein, in PtaeV_2 (GECLV1) and GbilV (GECLV2) (Fig. 3). CD searches did not find any integrase-like domain, a pattern similar to the angiosperm *Caulimoviridae* and indicates integration might not be necessary for the replication of gymnosperm EPRVs either.

**Figure 3.**
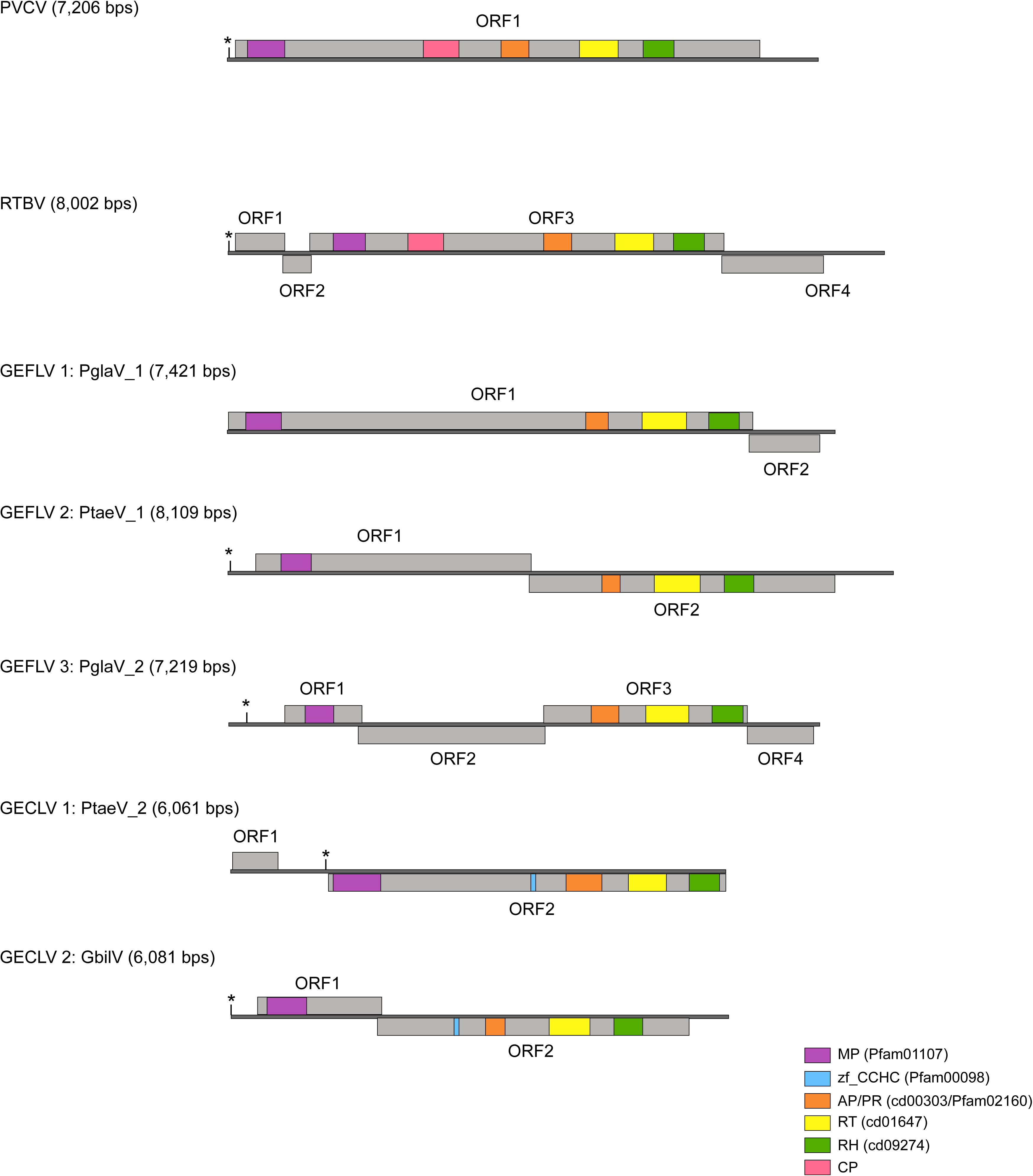
Genome structures of representative gymnosperm EPRVs and exogenous *Caulimoviridae*. Virus name abbreviation: RTBV, Rice tungro bacilliform virus; PVCV, Petunia vein clearing virus; PglaV_1, *Picea glauca* virus 1; PglaV_2, *Picea glauca* virus 2; PtaeV_1, *Pinus taeda* virus 1; PtaeV_2, *Pinus taeda* virus 2; GbilV, *Ginkgo biloba* virus. The tRNA^Met^ which represents the beginning of the viral replication, was indicated by an asterisk. The grey lines represent the genomes and the rectangles represent the putative open reading frames (ORFs). The conserved protein domains were labelled in different colors. Abbreviations: MP, movement protein; PR/AP: protease/pepsin-like aspartate protease; RT, reverse transcriptase; RH, ribonuclease H1; zf_CCHC, zinc-finger CCHC motif.

**Figure 4.**
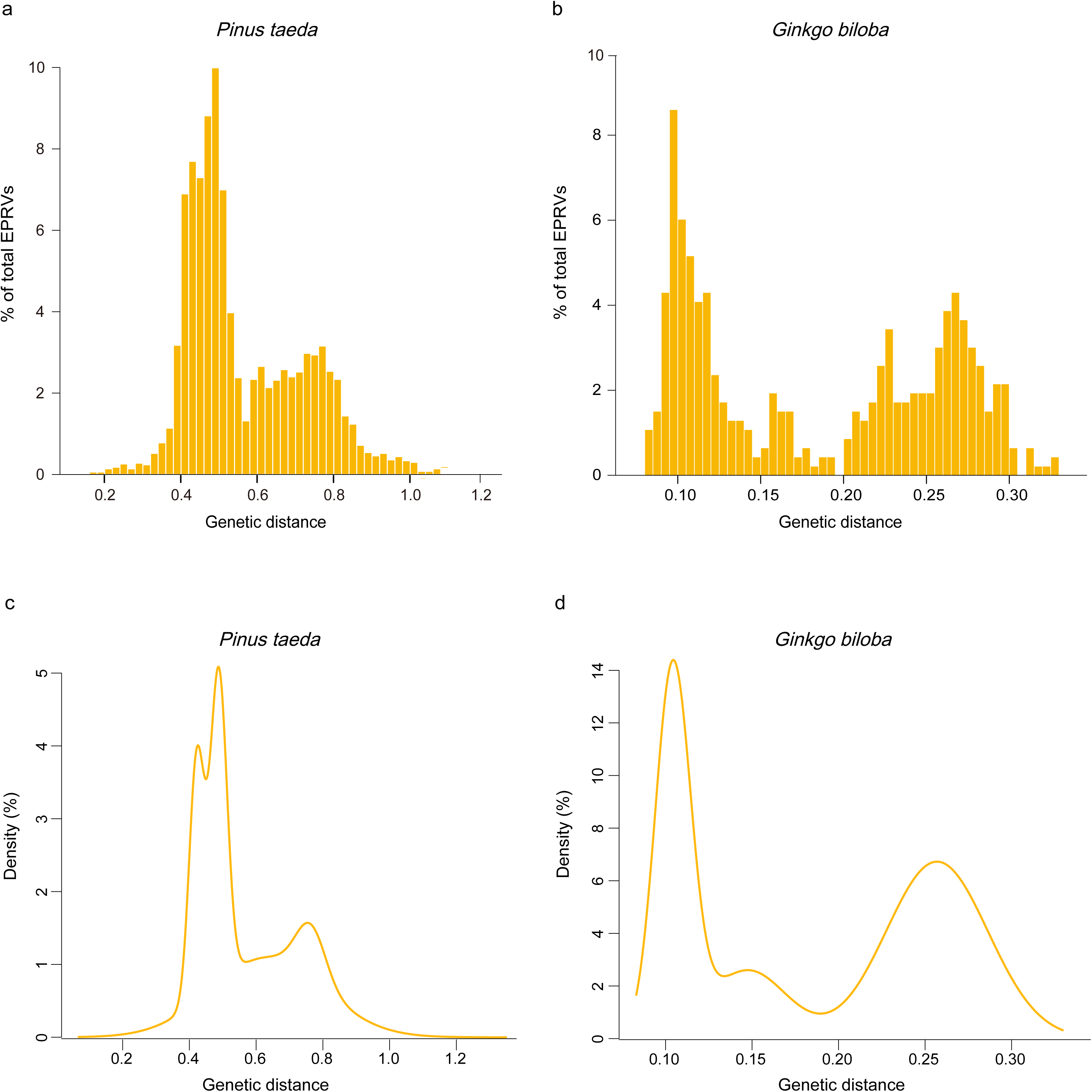
Proliferation dynamics of EPRVs within the *P. taeda* (a and c) and *G. biloba* (b and d) genomes. Yellow rectangles in **a** and **c** represent the distribution of the genetic distances between EPRV copies and their consensus sequences. **b** and **d** indicate Gaussian mixture models fitted.

### Age estimate of EPRV bursts

Because the genomes of loblolly pine (*P. taeda*) and ginkgo (*G. biloba*) were of relatively high quality and contain rather distinct numbers of EPRVs (Table 1), they were used to infer the evolutionary dynamics of EPRVs within the host genomes. Mixture model analyses of the genetic divergence between EPRV copies and their consensus nucleotide sequence show that there are four and three peaks in *P. taeda* and *G. biloba* (Supplemental Table 2), suggesting at least four and three independent EPRV integration events occurring along the lineages leading to *P. taeda* and *G. biloba*, respectively. Based on the mixture analyses and phylogenetic analyses (Supplemental Fig. 2), the ERPVs within the genomes of *P. taeda* and *G. biloba* were classified into four and three families.

We failed to find any orthologous integration of EPRVs in different species and cannot directly estimate the time of viral integration. The median pairwise genetic distance within each family was calculated to examine the age of burst for each EPRV family^22, 23^ (Supplemental Table 3). Our results show that the EPRV proliferation dynamics were of difference between *P. taeda* and *G. biloba* (Supplemental Table 3). But we found all of the EPRV families investigated here experienced proliferation peaks tens or hundreds of million years ago. Consistently, the EPRV copies contain many frame-shift mutations and premature stop codons (Supplemental Fig. 4). However, those analyses come with two caveats: i) it is uncertain whether the EPRV proliferation activity within the host genome follows the Gaussian distribution; ii) the evolutionary rate of EPRVs remains unclear. Nevertheless, our results suggest that EPRVs evolved within their host genomes for hundreds of millions of years and indicate an ancient origin of *Caulimoviridae*.

## Discussion

In this study, we report the identification of EPRVs within the genomes of gymnosperms and ferns. Together with the previous reports of exogenous and endogenous *Caulimoviridae* in angiosperms, our results demonstrate that all the major lineages of euphyllophytes (ferns and seed plants) are/were infected by the *Caulimoviridae* family. No EPRVs detected in the genome of *C. richardii* does not necessarily mean the absence of EPRVs, which is probably due to the low coverage nature of its genome assembly. Few viruses have been documented in plant species outside angiosperm^24^. The identification of EPRVs in gymnosperms and ferns makes *Caulimoviridae* the only known viral family that infects all major lineages of euphyllophytes.

Our findings show that the newly identified EPRVs exhibit an unprecedented diversity, and the known angiosperm viral diversity only accounts for a minority of the *Caulimoviridae* diversity. The current *Caulimoviridae* classification system^16^ cannot readily account for the diversity of EPRVs in gymnosperms and ferns. Indeed, the divergence within one clade of gymnosperm or fern EPRVs is comparable to the divergence of one exogenous viral genus or florendoviruses. Therefore, an updated classification incorporating gymnosperm and fern EPRVs should be developed. Most of the EPRV clades lack the exogenous counterparts, either because the ancient viral lineages completely died out, or because many exogenous viruses remain to be discovered.

Two possible macroevolutionary modes of *Caulimoviridae* could be conceived: i) Co-speciation model: the viruses have co-evolved with their euphyllophyte hosts for >400 million years and undergone sporadic cross-species transmission (Fig. 5a); ii) Cross-species transmission model: frequent cross-species transmissions predominated in the evolution of *Caulimoviridae* (Fig. 5b). In this study, we failed to find co-speciation signal between *Caulimoviridae* and its hosts, suggesting co-speciation might not be predominant in the macroevolution of *Caulimoviridae* (Table 2 and Supplemental Fig. 3). Indeed, it appears that the angiosperm viruses originated multiple times via independent cross-division transmission events (gymnosperms to angiosperms). For plant viruses, cross-division transmission events were rarely documented, partially because much remains unknown about the virosphere in plants outside angiosperms. Given multiple cross-division transmission events took place in *Caulimoviridae*, we think that cross-division transmission might not be a rare event for other plant viruses.

**Figure 5.**
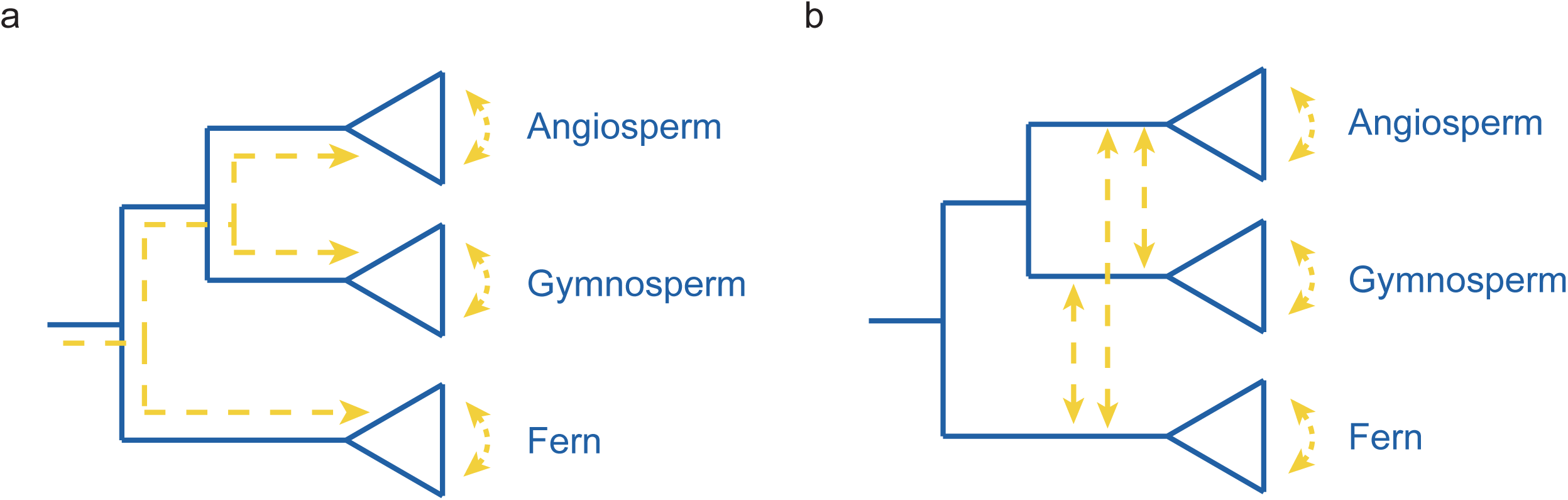
The models of the *Caulimoviridae* macroevolution. The evolution of plant hosts and viruses were indicated by blue lines and yellow dash lines, respectively. (**a**) Co-speciation model: the viruses have co-evolved with their euphyllophyte hosts and undergone sporadic cross-species transmission. (**b**) Cross-species transmission model: frequent cross-species transmission predominated in the evolution of *Caulimoviridae*.

Phylogenetic analysis and ancestral state reconstruction show that the *Caulimoviridae* family might have a monilophyte origin (Supplemental Fig. 1), which is compatible with the fact that we did not find any EPRV in earlier branching plants (lycophytes and nonvascular plants). One might argue that the absence of EPRVs in earlier branching plants is due to no viral integration occurring. However, this possibility seems to be unlikely, given viral integration is so widespread in euphyllophytes. The paleoviruses provide ‘molecular fossils’ for estimating the age of related viruses. Previously, the integration of banana streak virus into the *Musa balbisiana* genome was estimated to occur 0.63 million years ago^25, 26^. The endogenization of florendoviruses in *Oryza* speices was estimated to take place at least 1.8 million years ago^20^. Although we cannot directly date when the EPRV endogenization events occurred by finding orthologous integration of EPRVs in closely related species, the proliferation dynamics analyses of EPRVs within two representative genomes indicate they might have activated within the host genomes for hundreds of millions of years. Taken together, our findings pinpoint a possible ancient monilophyte origin of *Caulimoviridae*.

Unlike EVEs of non-retroviral source, the copy numbers of EPRVs are generally high (>1,000 copies for 10 out of 16 gymnosperm and fern species), suggesting that EPRVs contribute significantly to the complexity of host genomes. *Caulimoviridae* are closely related to LTR retrotransposons^11^. On the other hand, EPRVs lack LTRs, which makes it inappropriate to be classified as an LTR retrotransposon. It seems to be more appropriate to define EPRVs as a unique group of transposable elements^19^.

Our findings suggest that *Caulimoviridae* integrated into and amplified within host genomes multiple times. However, the integration and amplification mechanisms of EPRVs remain unclear, as the *Caulimoviridae* genomes lack integrase-like proteins and integration is not essential for their replication. Several potential mechanisms might be involved: i) Unlike the highly sequestered nature of animal germlines, plant germline tissues might allow more frequent infections of viruses. However, the copy numbers of EVEs within animals seem to be generally similar to EVEs within plants^5, 6^. It appears that the high copy numbers of EPRVs are due to its own biology. ii) EPRVs encode a ‘cryptic’ integrase without significant similarity with the known proteins that function in integration. No integrase domain was found in the *Petunia vein clearing virus* (PVCV) genome. But one of its proteins encodes two distinctive motifs [HHCC and DD(35)E] that are shared by the integrase domain of retroviruses and LTR retrotransposons^27^. However, it remains unknown whether that protein performs function similar to integrase. iii) Microhomology-mediated recombination between EPRV sequences and host sequences during host gap repair process could result in the integration of viral sequences into the host genomes^17^. This mechanism requires the free ends of open circular viral sequences produced during virus replication^19,28^. iv) Like short interspersed elements (SINE), EPRVs might integrate and amplify themselves within the host genomes via hijacking the integrase of other retrotransposons^29,30^.

## Method

### Identification of EPRVs in plant genomes

The genome sequences of twenty plant species were used to screen the presence of EPRVs, including ten gymnosperms (*P. taeda, Pinus lambertiana, Pinus sylvestris, Picea abies, Picea glauca, G. biloba, Gnetum gnemon, Juniperus communis, Taxus baccata*, and *Abies sibirica*), six ferns (*C. richardii, Dipteris conjugata, Plagiogyria formosana, Pteridium aquilinum, Polypodium glycyrrhiza*, and *Cystopteris protrusa*), one moss (*P. patens*), one liverwort (*M. polymorpha*), one lycophyte (*S. moellendorffii*), and one charophyte (*K. flaccidum*)^21,31,32^ (Table S1). To identify putative EPRVs within these genomes, we employed a two-step phylogenomic approach. First, the tBLASTn algorithm was employed to search against the plant genomes using the RT-RH domain sequences of *Rice tungro bacilliform virus* (RTBV) and PVCV as queries with an *e* cutoff value of 10^-10^. Next, all the significant hits obtained were aligned with RT-RH sequences of representative LTR retrotransposons, retroviruses, *Hepdnaviridae*, and *Caulimoviridae*^33^ using MAFFT with default parameters^34^. Putative EPRVs, which form a monophyletic group with other *Caulimoviridae* with high support values, were identified based on phylogenetic analyses. EPRVs were confirmed by further rounds of phylogenetic analyses with putative EPRVs and representative LTR retrotransposons, retroviruses, *Hepdnaviridae*, and *Caulimoviridae*. Phylogenetic analyses were performed using an approximate maximum likelihood method implemented in FastTree 2.1.9 with default parameters^35^. The copy number of EPRVs within each species was then counted. If length between hits was less than 5,000 bps and the hits were in the same order as the query, the hits were treated as a single copy.

### Phylogenetic analysis

To further analyze the relationship among *Caulimoviridae*, phylogenetic analysis was performed using the RT-RH protein sequences from representative EPRV sequences of each gymnosperm and fern species, exogenous viruses, and florendoviruses. The *Ty3* retrotransposon sequence was used as the outgroup. These protein sequences were aligned using MAFFT algorithm with an accurate method with the L-INS-i strategy^34^. Ambiguous regions within the alignment were removed using Gblocks 0.91b^36^. The phylogenetic analysis was performed using a Bayesian method implemented in MrBayes 3.2.6 (ref. 37). The RtRev substitution model was used. A total of 17,000,000 generations in four chains were run, sampling posterior trees every 100 generations. The first 25% of the posterior trees were discarded for further analysis.

### Reconstruction of consensus genome sequences

Relatively complete EPRV sequences with *MP-AP-RT-RH* domains with extended flanking regions (∼5,000 bps for each end) were extracted. These sequences were then used as queries to search sequences with high similarity within their own host genome using the BLASTn algorithm with an *e* cutoff value of 10^-25^. The significant hits were aligned using MAFFT^34^ and consensus sequences were generated using Geneious 10 (ref. 38) and manually edited. ORFs with the minimum size of 500 were found using Geneious 10 (ref. 38). Protein domains within these reconstructed genomes were detected using the CD search^39^.

### Analysis of EPRV activity within host genomes

Because the genomes of *P. taeda* and *G. biloba* were of relatively high quality, we used their genomes to infer the evolutionary dynamics of EPRVs within the host genomes. The *AP-RT-RH* nucleotide sequences with length >500 bps of all the EPRVs within *P. taeda* and *G. biloba* genomes were extracted independently. These sequences were aligned using MAFFT and consensus sequences were inferred using Geneious 10 (ref. 38). The genetic distance between the consensus sequences and EPRVs was calculated based on the Kimura two-parameter model. To identify significant peaks in the genetic distance data sets, Gaussian mixture models were fitted using the R package mclust. The number of components (each component is modeled by the Gaussian distribution) was estimated by fitting models. The Bayesian Information Criterion (BIC) was used as the model selection criterion.

The phylogenetic trees of the *AP-RT-RH* nucleotide sequences extracted above within each host genome were reconstructed using FastTree 2.1.9 with a GTR＋CAT model. The different EPRV families within the phylogenetic trees were allocated based on the results by mixture model analyses. The nucleotide sequences of each EPRV family were extracted and aligned using MAFFT^34^. Pairwise genetic distances were calculated based on the Kimura two-parameter model. The age of burst (*T*) for each EPRV family was estimated through *T* = *D*/2*μ*, where *D* represents the median pairwise distance and *μ* represents the evolutionary rate of host (∼1.43×10^-9^-2.2×10^-9^ substitutions per site per year^31^).

### Co-speciation analysis

We explore the host-virus co-speciation signal at the level of class, for the complex evolutionary history of EPRVs after integration might complicate co-speciation analysis. The relationships between virus and host phylogenetic trees were assessed using an event-based method implemented in Jane 4 (ref. 40). Briefly, five events (cospeciation, duplication, duplication & host switch, loss and failure to diverge) were assigned to a cost. The number of each event was estimated by finding the solution with the minimum total cost. The event-cost schemes (cospeciation-duplication-duplication & host switch-loss-failure to diverge) were set as follows, -1-0-0-0-0 (refs. 41, 42) and 0-1-1-2-0 (ref. 40). Host-virus phylogeny congruence was assessed by statistical tests with both two randomization methods, random tip mapping and random parasite tree, with the sample size of 500.

### Reconstruction of ancestral states

To detect the macroevolutionary pattern among *Caulimoviridae*, we performed ancestral state reconstruction with Mesquite 3.10 (ref. 43). We assigned the 96 viral taxa (Fig. 2) using their hosts (gymnosperm, angiosperm, and fern) as characters. For the outgroup Ty3, a ‘?’ character was assigned. The parsimony model was used to trace character evolution over the posterior trees sampled in the aforementioned Bayesian analysis.

## Acknowledgements

This work was supported by the Natural Science Foundation of Jiangsu Province (BK20161016) and the Priority Academic Program Development (PAPD) of Jiangsu Higher Education Institutions.

## Author contributions

G.-Z.H. designed the study. G.Z. and G.-Z.H performed the experiments. G.-Z.H and G.Z. analyzed the data and wrote the manuscript.

